# Divalent metal cations potentiate the predatory capacity of amoeba for *Cryptococcus neoformans*

**DOI:** 10.1101/214460

**Authors:** Man Shun Fu, Arturo Casadevall

## Abstract

Among the best studied interaction between soil phagocytic predators and a human pathogenic fungus is that of *Acanthamoeba castellanii* and *Cryptococcus neoformans*. The experimental conditions used in amoeba-fungal confrontation assays can have major effects on whether the fungus or the protozoan is ascendant in the interaction. In the presence of Mg^2+^ and Ca^2+^ in PBS, *C. neoformans* was consistently killed when incubated with *A. castellanii*. *A. castellanii* survived better in the presence of Mg^2+^ and Ca^2+^, even when incubated with *C. neoformans*. In the absence of Mg^2+^ and Ca^2+^, *C. neoformans* survived when incubated with *A. castellanii*, and the percentage of dead amoeba was higher than when incubated without yeast cells. These results show that the presence of Mg^2+^ and Ca^2+^ can make a decisive contribution toward tilting the outcome of the interaction in favor of amoeba. Of the two metals Mg^2+^ had a stronger effect than Ca^2+^. Cations enhanced *A. castellanii* activity against *C. neoformans* through enhanced phagocytosis, which is the major mechanism for amoeba to kill fungal cells. We found no evidence that amoeba uses extracellular killing mechanisms in their interactions with *C. neoformans*. In summary, the presence of Mg^2+^ and Ca^2+^ enhanced cell adhesion on surface and motility of amoeba, thus increasing the chance for contact of *C. neoformans* and the frequency of phagocytosis. Our findings imply that divalent cation concentration in soils could be an important variable for whether amoeba can control *C. neoformans* in the environment.

**Importance:** Grazing of soil organisms by phagocytic predators such as amoeba is thought to select for traits that allow some of them to acquire the capacity for virulence in animals. Consequently, knowledge about the interactions between amoeba and soil microbes, such as pathogenic fungi, is important for understanding how virulence can emerge. We show that the interaction between amoeba and the pathogenic fungus *C. neoformans* is influenced by the presence of magnesium and calcium in the assay, which potentiate amoeba. The results may also have practical applications since enriching soils with divalent cations may reduce *C. neoformans* numbers in contaminated soils.

## Introduction

*Cryptococcus neoformans* is a soil-dwelling fungus that is a frequent cause of life-threatening meningoencephalitis in individuals with impaired immunity (1). One of the fascinating aspects of *C. neoformans* biology is that it has the capacity for virulence in very different animal and plant species yet this organism has no need for pathogenicity, as it can survive in soils without hosts. To explain this phenomenon, the concept of accidental virulence was proposed, which posits that environmental pressures select for traits that confer upon the microbe the capacity for pathogenicity independent of the final host (2, 3). In the case of *C. neoformans* the environmental pressure was proposed to include amoeboid predators such as amoeba (4, 5). Supporting this hypothesis is the fact that many virulence factors function in the same manner during the interaction between macrophages and amoeba (5, 6). Furthermore, passage of an avirulent strain of *C. neoformans* in social amoeba restored virulence (7). Recently, the interaction of *C. neoformans* with *A. castellanii* was shown to result in increased fungal virulence for Galleria mellonella as a result of amoeba-induced changes in susceptibility to microbicidal oxidants and cell wall (8). Studies of amoeba interactions with other pathogenic fungi such as *Histoplasma capsulatum* (9), *Sporothrix schenkii* (9), *Blastomyces dermatitides* (9), *Aspergillus fumigatus* (10, 11), and entomopathogenic fungi (12) suggest that fungal-amoeba interactions could be important in their virulence for animal hosts.

In recent years amoeba have emerged as a major system for studying bacterial and fungal host-microbe interactions (13, 14), including *C. neoformans* biology and virulence (15-19). The realization that amoeba served as reservoirs and training hosts for animal virulence together with the ease that these protozoa can be maintained in the laboratory have popularized the study of their interactions with various microbes. Insights gained with amoeba on mechanisms of intracellular pathogenesis were shown to apply to microbe-macrophage interactions and vice versa. For example, the discovery that phospholipids are a trigger for capsular enlargement in *C. neoformans* followed the observation that cryptococcal cells enlarged their capsules in the presence of amoeba, and then the same phenomenon was shown in macrophages (20). On the other hand, a *Mycobacterium avium* pathogenicity island important for intracellular infection was first identified in a macrophage screen and then shown to be also important for infecting amoeba (21). Clearly, macrophages and amoeba provide complementary systems for the study of virulence determinants and the evolution of pathogenicity.

Prior studies have shown that the outcome of the *C. neoformans-Acanthamoeba castellanii* interaction is highly dependent on the conditions of the experiment. For example, confrontation experiments involving cryptococci and amoeba in nutrient poor conditions such as phosphate buffered saline resulted in fungal growth and death of amoeba (4, 22). However, when cryptococci and amoeba were suspended on amoeba growth media the protozoa were ascendant, with reduction of fungal cells by predation and killing (23). Those results were interpreted as suggesting that the nutritional state of amoeba was an important variable in their predatory capacity (23). However, while carrying out *C. neoformans*-amoeba experiments we made the serendipitous observation that the presence of Ca^2+^ and Mg^2+^ was all that was required to potentiate amoeba predation for fungal cells. These results identify the concentration of divalent cations as an important variable in studies of microbe-amoeba interactions.

## Results

### A serendipitous observation

While studying *C. neoformans* interactions with *A. castellanii* in phosphate buffered saline we noted that the amoeba killed a substantial proportion of the fungal cells. This result was unexpected given that prior work has shown that in these conditions *C. neoformans* tended to kill *A. castellanii* and replicate (4, 23). A review of the conditions of the experiment revealed that the solution used was not the standard laboratory formulation but instead a commercial product manufactured by Corning (Corning, NY) known as Dulbecco’s Phosphate Buffered Saline, which differed from the PBS used in prior experiments by supplementation with magnesium and calcium. This suggested that the difference in the findings was the presence of these two divalent cations. We then repeated the experiments with DPBS and supplemented with and without Ca^2+^ and Mg^2+^, jointly and singly, and verified that addition of the metal salts was the ingredient potentiating the activity of *A. castellanii* against *C. neoformans*. In the presence of magnesium and calcium in DPBS, *C. neoformans* CFUs decreased when incubated with *A. castellanii* but increased without *A. castellanii* (Figure 1A and B). *A. castellanii* survived better in the presence compared to the absence of Ca^2+^ and Mg^2+^ during the incubation with *C. neoformans* (Figure 1C). The percentage of dead *A. castellanii* after incubation with *C. neoformans* was higher than without yeast cells in the absence of divalent metal cations (Figure 1C). The results show that the presence of magnesium and calcium are key determinants of the outcome of *C. neoformans* and amoeba interaction.

**Figure 1.**
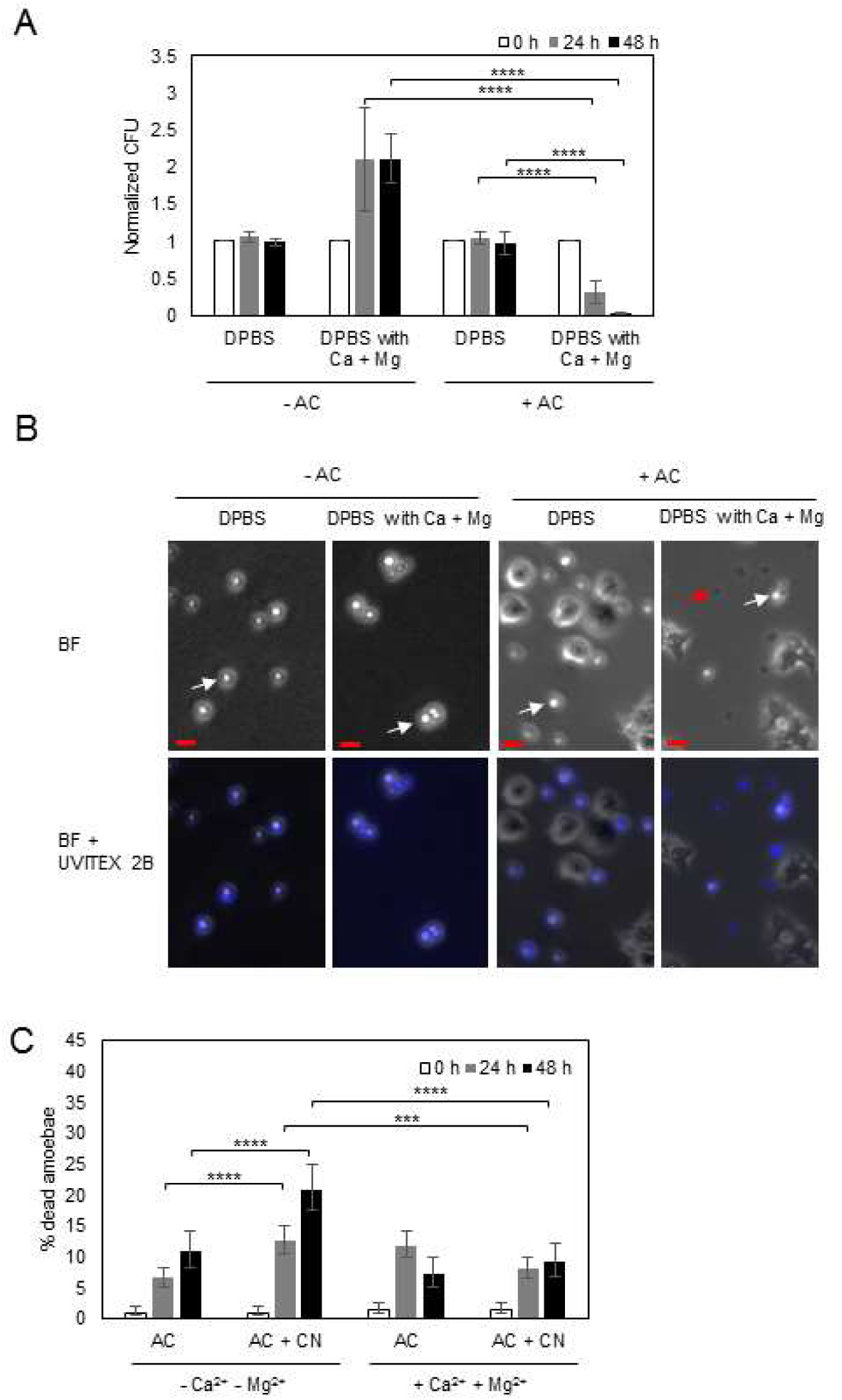
The presence of magnesium and calcium affects the outcome of the *A. castellanii*-*C. neoformans* interaction. (A) The presence of magnesium and calcium decreases the survival of *C. neoformans* (CN) during the incubation with *A. castellanii* (AC). The survival of *C. neoformans* was determined by CFU after incubation with *A. castellanii* for 0, 24 and 48 h. The CFU counts at 24 h and 48 h were normalized to the initial CFU at time zero. Data represent the mean of four biological samples. Error bars are SD. **** P < 0.0001 by Student’s t test. (B) Phase contrast and fluorescence images of Uvitex 2B stained *C. neoformans* co-incubated with or without *A. castellanii* in DPBS with or without magnesium and calcium. In the phase contrast images, disrupted *C. neoformans* are dark (red arrow) while intact cells are refractile (white arrow). More disrupted *C. neoformans* appeared in the condition with *A. castellanii* incubation in DPBS containing magnesium and calcium. Scale bar is 10 µm. (C) *A. castellanii* survived better in the presence of Mg^2+^ and Ca^2+^ during the incubation with *C. neoformans*. The viability of *A. castellanii* were determined by trypan blue exclusion assay. The percentage of dead *A. castellanii* is determine by counting the number of blue staining cells per total cell number counted. Four independent biological experiments were performed. Error bars represent 95 % confidence interval of the mean. ***P = 0.0008 **** P < 0.0001 by Fisher’s exact test.

The commercial DPBS buffer supplemented with CaCl_2_ and MgCl_2_. DPBS (without Ca^2+^ and Mg^2+^) also contains KCl and NaCl, ruling out chloride ions as the component responsible for the enhanced predation of amoeba. However, to further confirm that the divalent metals were responsible for the enhanced protozoal fungicidal effects, we performed the experiment with DPBS supplemented with different forms of Ca^2+^ and Mg^2+^ (calcium nitrate and magnesium sulfate). In these conditions we observed enhanced killing ability of amoeba, establishing that magnesium and calcium ions are the ones responsible for this effect (Fig S1).

### Mg^2+^ is more effective than Ca^2+^ in potentiating *A. castellanii*

To ascertain which of the divalent cations was responsible for potentiating amoeba we evaluated the outcome of the *C. neoformans*-*A. castellanii* interaction in conditions where DPBS was supplemented with Ca^2+^, Mg^2+^ or both. Addition of either Ca^2+^ or Mg^2+^ had a significant effect in reducing *C. neoformans* CFU relative to conditions with no divalent cations but the effect was much greater with Mg^2+^, and maximal when both cations were present simultaneously (Figure 2A-D). The presence of Mg^2+^ also increased the viability of *A. castellanii* in the presence and absence of *C. neoformans* (Figure 2G).

**Figure 2.**
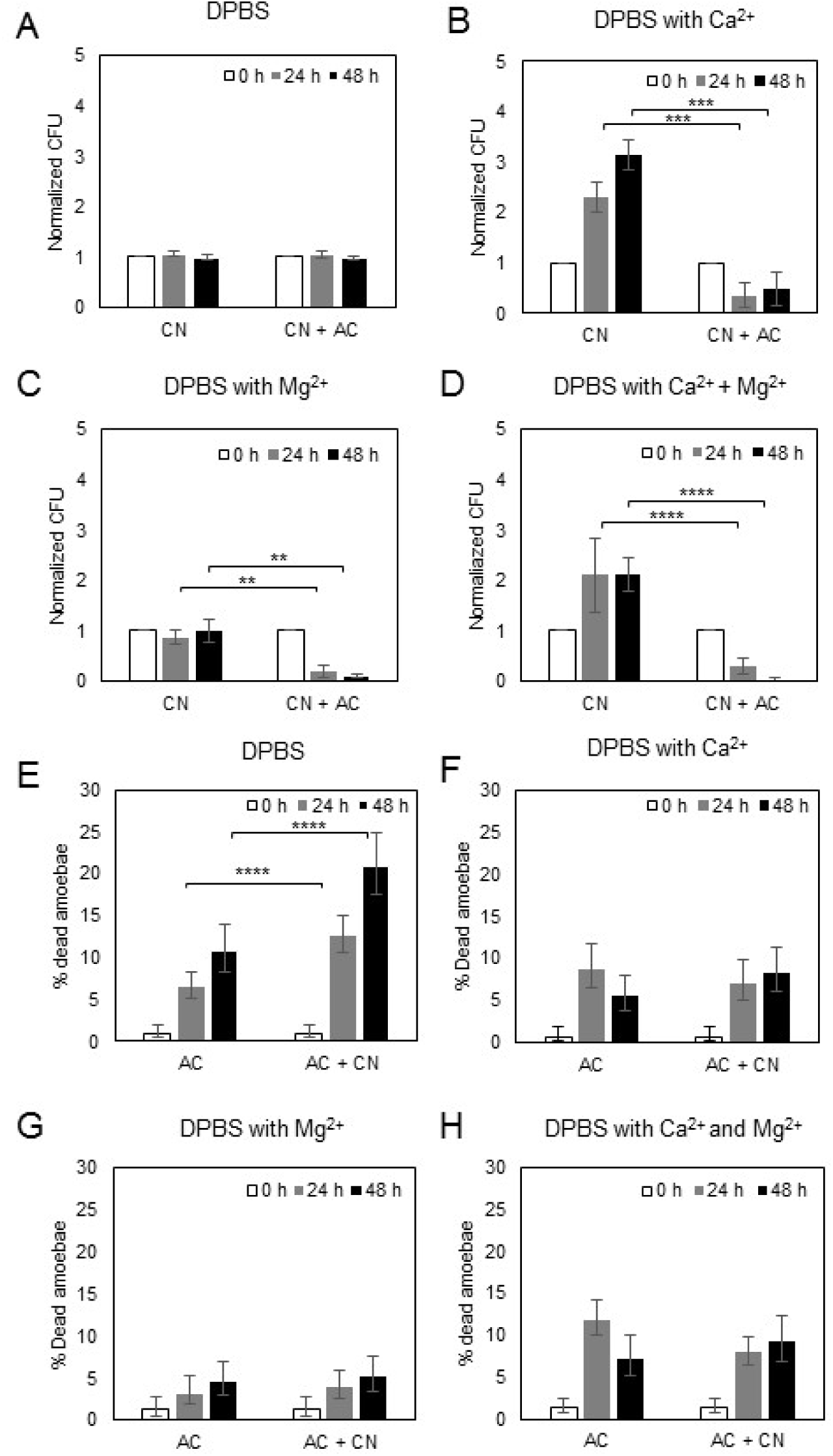
Magnesium had a stronger effect than calcium on reducing the survival of *C. neoformans* when incubated with *A. castellanii*. The survival of *C. neoformans* (CN) were determined by CFU after incubation with *A. castellanii* (AC) for 0, 24 and 48 h in (A) DBPS, (B) DPBS with calcium, (C) DPBS with magnesium, and (D) DPBS with calcium and magnesium. The CFU counts were normalized to the initial CFU at time zero. Data represent the mean of three biological samples. Error bars are SD. ** P < 0.01*** P < 0.001 ****P < 0.0001 by Student’s t test. The viability of *A. castellanii* were also determined by trypan blue exclusion assay after incubation with *C. neoformans* in (E) DBPS, (F) DPBS with calcium, (G) DPBS with magnesium, and (H) DPBS with calcium and magnesium. The percentage of dead *A. castellanii* is determine by counting the number of blue staining cells per total cell number counted. Three independent biological experiments were performed. Error bars represent 95 % confidence interval of the mean. **** P < 0.0001 by Fisher’s exact test.

### Divalent cations enhanced amoeba surface area, phagocytosis and mobility

To investigate the mechanism by which divalent cations potentiated *A. castellanii* against *C. neoformans* we evaluated three parameters with a high likelihood to impact the fungal-protozoal interaction: amoeba adhesion, phagocytosis and mobility. Both Ca^2+^ and Mg^2+^ are known to be important for cellular adhesion, with Mg^2+^ having a stronger effect than Ca^2+^ (24, 25). We measured cellular area as a measure of adhesion and noted that amoeba in the presence of Ca^2+^ and Mg^2+^ manifested significantly greater areas when placed on a glass surface (Figure 3A and B). Amoeba moved more and made more contacts with *C. neoformans* in the presence of Ca^2+^ and Mg^2+^ (Figure 3C and D), which translated into significantly greater phagocytosis of yeast cells (Figure 3E).

**Figure 3.**
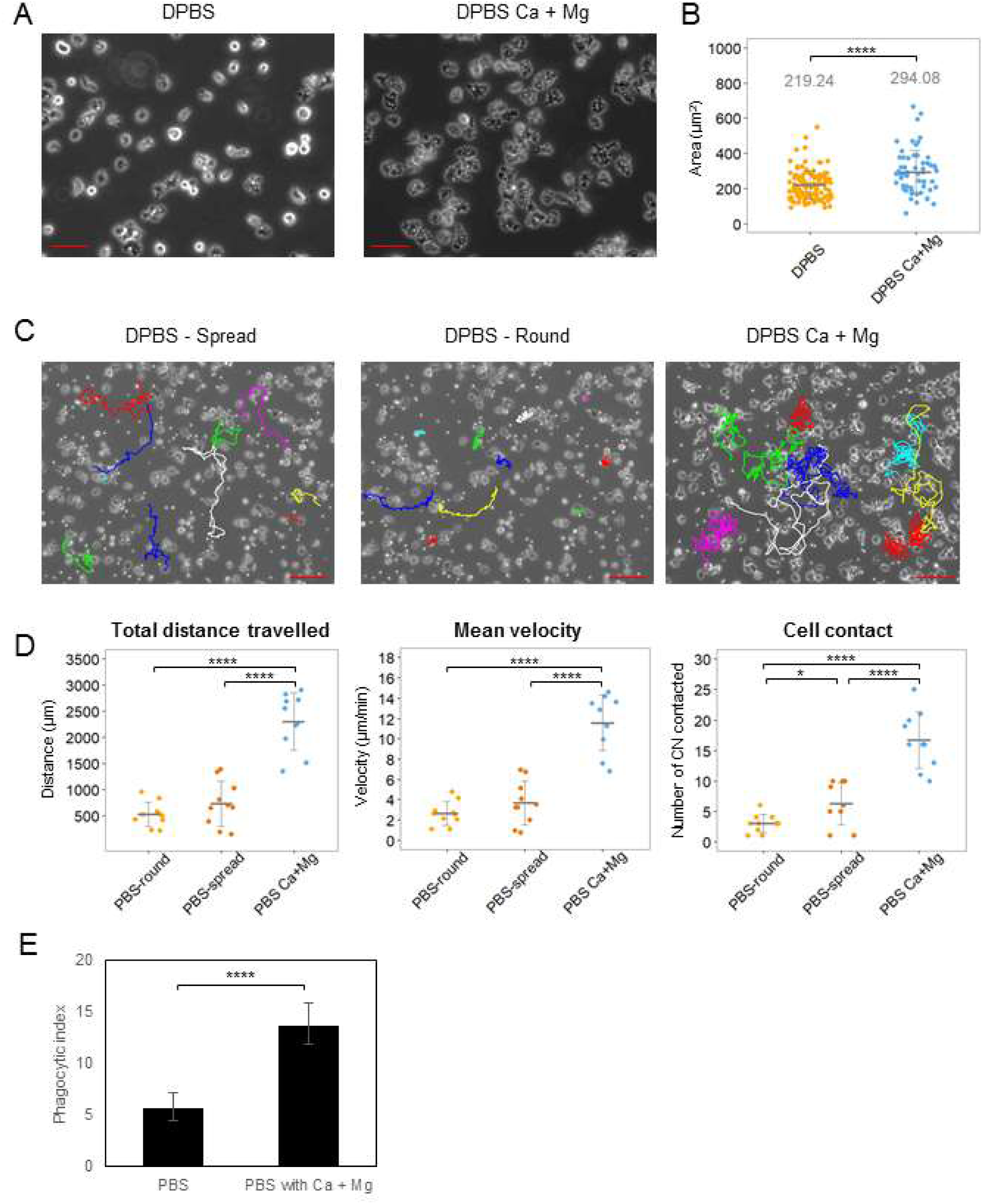
The presence of calcium and magnesium enhances the adherence of *A. castellanii* to surface and its motility. (A) Phase-contrast image of *A. castellanii* on glass surface in DPBS with or without Ca^2+^ and Mg^2+^. Scale bar is 50 µm. (B) To quantify cell spreading, the area of *A. castellanii* DPBS with or without Ca^2+^ and Mg^2+^ was measured. At least 100 cells were analyzed for each condition. Error bars indicate SD. **** P < 0.0001 by Student’s t-test. (C) Images of *A. castellanii* cell trajectories on glass surface in DPBS with or without Ca^2+^ and Mg^2+^. Ten *A. castellanii* cells were randomly selected for manually centroid tracked for total length of 2 h in each condition. The interval of each track is 30 s. Color lines indicate manual generated tracks. Two distinct morphologies of amoeboid cells, round and spread, in DPBS were tracked separately. Scale bar is 100 µm. (D) The total distance, the mean velocity and the frequency of contact with *C. neoformans* were qualified. The total distance was defined as the sum of distance the amoeboid cells travelled from starting point to end point of cell trajectory. The Mean velocity was calculated as the mean of all velocity measurements from amoeboid cells moving in each 30 s interval. Error bars indicate SD. *** P < 0.001, **** P < 0.0001 by Student’s t-test. (E) The presence of magnesium and calcium induces the phagocytosis of *C. neoformans* by *A. castellanii*. *A. castellanii* and Uvitex 2B labeled *C. neoformans* were incubated at 1:1 ratio in DPBS with or without magnesium and calcium for 2 h to allow phagocytosis. The phagocytic index was determined by the number of internalized *C. neoformans* per 100 *A. castellanii*. Data was obtained from four biologically independent experiments. Error bars represent 95 % confidence interval of the mean. **** P < 0.0001 by Fisher’s exact test.

### Amoeba fungicidal activity requires protozoa-fungal cell contact

Next, we considered whether contact was necessary for amoeba to kill *C. neoformans* by separating fungal and amoeba cells in wells where fluid was connected through a semipermeable membrane. We observed no reduction in fungal CFUs in conditions where the fungal and protozoal cells were separated. However, when *C. neoformans* and amoeba were placed in the same chamber there was a reduction in fungal colonies (Figure 4A and B). Incubation of *C. neoformans* in amoeba conditioned media, which is collected after co-incubation of *A. castellanii* and *C. neoformans* had no effect on fungal colonies arguing against the release of fungicidal products from amoeba (Figure 4C, 4D and Movie S1). These experiments indicated a necessity for contact between amoeba and *C. neoformans* for protozoal fungicidal activity. Cinematographic analysis of the *C. neoformans*-*A. castellanii* interaction revealed phagocytosis followed by regurgitation of fungal carcasses, with the latter being easily distinguishable from live fungal cells as these were shriveled structures that had lost their characteristic translucent appearance by light microscopy (Figure 4E). The average time from phagocytosis to regurgitation of cryptococcal cellular remnants was 2.04 ± 0.53 h (n = 5).

**Figure 4.**
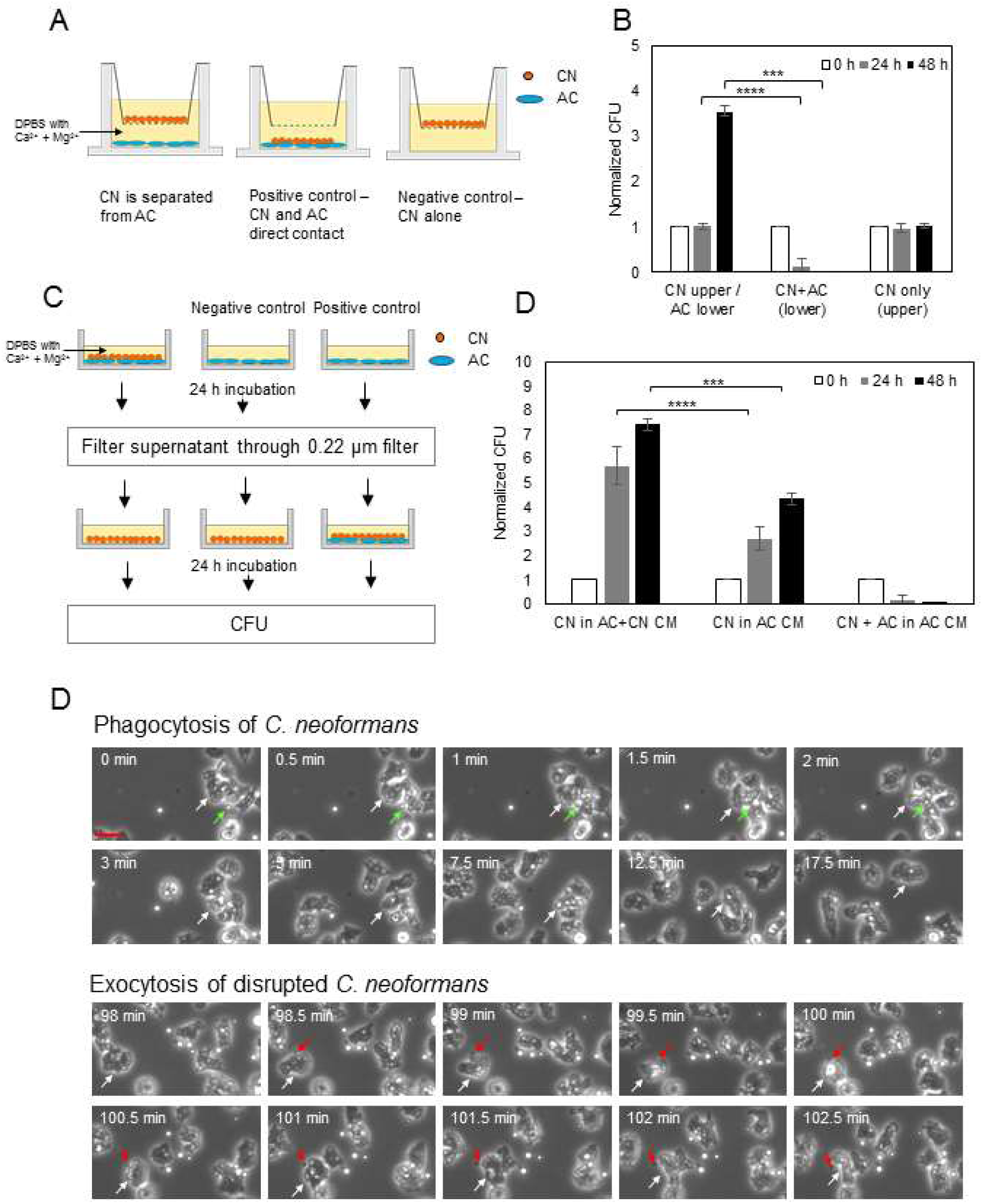
Phagocytosis is the major mechanism for *A. castellanii* to kill *C. neoformans* (A) Scheme of the Transwell^®^ assay. *A. castellanii* and *C. neoformans* were separated in DPBS with Ca^2+^ and Mg^2+^ by porous membrane (pore size 0.4 µm). The upper compartment contained *C. neoformans* while the lower compartment contained *A. castellanii*. As a positive control *C. neoformans* was incubated with *A. castellanii* at the lower compartment. As a negative control *C. neoformans* was incubated without *A. castellanii* at the upper compartment. (B) The survival of *C. neoformans* were determined by CFU after the Transwell^®^ assay for 0, 24 and 48 h. The CFU counts were normalized to the initial CFU at time zero. Data represent the mean of four biological samples. Error bars are SD. *** P = 0.0004, ****<0.0001 by Student’s t test. (C) Schematic for the production of conditioned media from co-incubation of *C. neoformans* (CN) and *A. castellanii* (AC) and exposure of the fungal cell. After exposure of *A. castellanii* to *C. neoformans* in DPBS with Ca^2+^ and Mg^2+^, conditioned media were collected, filtered, and added to fresh culture of *C. neoformans*. As a negative control *A. castellanii* conditioned medium was collected and added to *C. neoformans*. As a positive control *A. castellanii* conditioned medium was collected and added to the co-incubation of *C. neoformans* and *A. castellanii*. (D) The survival of *C. neoformans* (CN) were determined by CFU after exposure of conditioned medium (CM) for 0, 24 and 48 h. The CFU counts were normalized to the initial CFU at time zero. Data represent the mean of three biological samples. Error bars are SD. *** P = 0.00223 **** P < 0.0001 by Student’s t test. (E) Time-lapse phase contrast images of co-incubation of *C. neoformans* and *A. castellanii* in DPBS with Ca^2+^ and Mg^2+^. Upper panel exhibits the phagocytosis of *C. neoformans* (green arrow) by *A. castellanii* (white arrow). Lower panel shows the exocytosis of disrupted *C. neoformans* (appears in dark; red arrow) from *A. castellanii* (white arrow). Scale bar is 25 µm.

### Interaction of *C. neoformans* and amoeba

To better understand how divalent cations potentiated amoeba against *C. neoformans* we carried out several experiments to probe variables that might contribute to their interaction. Given that the capsule of *C. neoformans* has negatively charged glucuronic acid residues that can interact with cations and that capsule enlargement can be triggered by Ca^2+^ (26), we first incubated yeast cells with Mg^2+^ and Ca^2+^, washed and then added the fungal cells to amoeba (Figure 5A-B). We measured no effect suggesting that the increased susceptibility of *C. neoformans* to amoeba in the presence of cations was not due to effects on the yeast cell and/or capsule. Pre-treatment of the amoeba or both amoeba and *C. neoformans* with Mg^2+^ and Ca^2+^ (Figure 4C-F) followed by washing and incubation with *C. neoformans* also had no effect on fungicidal activity. These results indicate that the divalent cations have to be in the solution containing amoeba and fungal cells for their potentiating effect on amoeba fungicidal activity.

## Discussion

Amoeba are an increasingly popular alternative host system for studying interactions between microbes and environmental phagocytic cells but relatively little work has been done on the effect of experimental conditions on the outcome of their interaction with fungi. In earlier studies we had noted that carrying out confrontation experiments between *A. castellanii* and *C. neoformans* in ATCC Medium 712 PYG potentiated the fungicidal capacity of the protozoa and inferred that it reflected improved nutritional status for the amoeboid cells (23). Here we show that addition of the divalent cations Ca^2+^ and Mg^2+^ is sufficient to potentiate *A. castellanii* activity against *C. neoformans* and making it very likely that they are the active ingredients in the ATCC Medium 712 medium, which has 0.05 M CaCl_2_ and 0.4 M MgSO_4_. Our findings establish that the ionic composition of the solution can have an important effect on the outcome of the experiment with *C. neoformans* and suggest that the same may apply in other amoeba-microbial confrontation systems.

The ability of amoeba to pray on *C. neoformans* was first described in the 1970s (27, 28). Evidence for the importance of this interaction in the environment comes from the observation that the prevalence of *C. neoformans* in soils is reduced proportionally to the presence of amoeba (29). Consequently, amoeba probably play a major role in reducing fungal burden in contaminated soils and, as such, may reduce the amount of human and animal exposure. Our results suggest that the efficacy of amoeba in soils may depend on environmental divalent cation concentrations. In this regard, soils can vary greatly on free Ca and Mg depending on the presence of vegetable matter detritus and pH, which in turn can affect worm concentrations (30). In the United States soils Ca and Mg concentrations average 0.375 M (range 0.002 to > 4 M) and 0.6 M (range 0.0025 to 8 M), respectively (31). Such range in concentrations imply that the availability of these cations to soil amoeba would vary greatly depending on the soils given that that these concentrations do not necessarily imply that the element would be available in soluble cationic form.

The potentiation of amoeba fungicidal activity by divalent cation ions was associated with increased surface area, mobility, contact with fungal cells and phagocytosis. Hence, divalent cations appear to have global effects of amoeba cell biological function that may work synergistically to enhance their predatory capacity. Increased surface area together with greater mobility and enhanced contacts with *C. neoformans* cells are likely to increase the probability of phagocytosis, which was indeed measured. Amoeba were visualized ingesting cryptococcal cells that were later regurgitated as opaque and shriveled forms that were easily distinguishable from live translucent cells. The requirement for contact and phagocytosis in fungal killing was shown by the absence of fungicidal activity through diffusible substances. In contrast, facilitating contact between amoeba and yeast cells reduced fungal CFUs. Treatment of fungal cells with divalent cations had no discernible impact on the interaction suggesting that the effect of Ca^2+^ and Mg^2+^ in augmenting amoeba fungicidal activity was primarily, if not exclusively, due to effects on the protozoan cells.

Although the mechanism by which Ca^2+^ and Mg^2+^ affect *A. castellanii* has not been investigated there is evidence from related systems that these cations can have powerful effects on amoeba. Pinocytosis and locomotion of amoeba can be influenced by monovalent and divalent cations (32). *A. castellanii* is known to accumulate Ca^2+^ in the presence of excess cation with cells forming electron dense deposits (33). Displacement of cell surface associated Ca^2+^ with drugs was shown to inhibit phagocytosis (34). Both Ca^2+^ and Mg^2+^ trigger signaling pathways that promote chemotaxis in the social amoeba Dictyostelium discoideum (35). For Entamoeba invadens the signaling program that triggers encystation is dependent on Ca^2+^ (36). Ca^2+^ has been shown to increase the adhesion of Acanthamoeba polyphaga to extracellular matrix proteins (37), in a manner that recapitulates the increased adhesion effects we describe in this study. However, not all interactions of amoeba with other cells require divalent cations. In this regard, *Entamoeba histolytica* adhered and killed Chinese Hamster Ovary cells in the presence of EDTA suggesting the divalent cations were not needed for the interaction (38). These observations illustrate the protean effects of divalent cations on protozoa and in aggregate support the view for major differences in the physiological state of amoeba cells in conditions with and without Ca^2+^ and Mg^2+^, which translate into large differences in their ability to kill *C. neoformans*. However, it is noteworthy that *C. neoformans* also responds to divalent cations with capsular enlargement (26), which protects against amoeba (39). Hence, the fungal-protozoal balance of power could reflect not only cation concentrations but also the time each type of organism had to adapt to the prevailing conditions.

Review of several publications describing amoeba interactions with various pathogens report different experimental conditions that could vary in divalent cations. For example, *A castellanii* interactions with *Vibrio cholerae* (40), *Aspergillus fumigatus* (10), *Pseudomonas aeruginosa* (41), *Corynebacterium spp*. (41) and *E. coli* (42), were studied in peptone yeast extract and glucose (PYG) media while interactions with *Campylobacter jejuni* (43), were done in ‘amoeba buffer’ that included Ca^2+^ and Mg^2+^ cations and *Bacillus anthracis* (44) in autoclaved creek water supplemented with divalent cations. Given the powerful effects of divalent cations on amoeba function comparisons across studies need to take into account the possibility that differences in Ca^2+^ and Mg^2+^ could affect experimental outcomes.

*C. neoformans* is commonly found in soils enriched in bird feces (reviewed in (1)). In urban centers *C. neoformans* is often found in high densities in soils near sites where pigeons roost where guano provides a rich source of nutrients (45). Human and animal infection is believed to originate from inhalation of desiccated yeast cells or basidiospores that become aerosolized from *C. neoformans* contaminated soils (1, 46). Consistent with an environmental source of infection isolates recovered from environmental sites were indistinguishable from those recovered from patients using several genetic markers (47, 48). Contaminated soils near a hospital have been suggested as sites for hospital-acquired infection of *C. neoformans* (49). Remediation of *C. neoformans*-contaminated soils is a formidable challenge and current methods involve the application the noxious chemicals such as formalin (50) and quaternary ammonium salts (51). Given that amoeba predation can reduce fungal burden in contaminated soils (29), our results suggest that increasing the soil concentration of divalent cations with calcium and magnesium salts could potentiate the fungicidal capacity of amoeba and provide a more environmentally friendly biological alternative to chemical decontamination.

In summary, we establish the divalent cation concentration is a major variable in the outcome of *A. castellanii* interactions with *C. neoformans*. Given the dramatic effects that divalent cations have on amoeba physiology we suspect that this effect will apply to other microbial-amoeba interactions and is likely to be an important experimental variable in other systems. This effect may also have applications for the environmental control of *C. neoformans* since increasing Ca and Mg concentration in soils could potentiate amoeba predation and reduce the fungal burden in contaminated sites.

## Materials and Methods

### Cell culture

Acanthamoeba *castellanii* strain 30234 was obtained from the American Type Culture Collection (ATCC). Culture was maintained in peptone-yeast extract-glucose (PYG) broth (ATCC medium 712) at 25 °C according to instructions from ATCC. *C. neoformans* var. *grubii* serotype A strain H99 was used in all experiments and this strain was originally obtained from Dr. John Perfect (Durham, NC). Cryptococcal cells were cultivated in Sabouraud dextrose broth with shaking (120 rpm) at 30°C for overnight (16 h) prior to use in amoeba assays.

### Assay of *A. castellanii* and *C. neoformans* interaction

The survival of *C. neoformans* in amoeba culture was performed as described previously (22). Briefly, *A. castellanii* were washed twice with Dulbecco's Phosphate-Buffered Saline (DPBS), Corning^®^ (Corning, Corning, NY) and diluted in DPBS to appropriate density. This buffer is available with and without calcium and magnesium supplementation. *A. castellanii* cells (1 × 10^4^ cells/well) were added to 96-well plates and allowed to adhere for 1 h at 25 °C. *C. neoformans* cells were washed twice with DPBS and diluted in DPBS to appropriate density. Fungal cells (1 × 10^4^) were added to wells containing amoeba or control wells containing DPBS alone, and the plates were incubated at 25 °C. At 0, 24, and 48 h, the amoeba were lysed by pulling the culture through a 27-gauge syringe needles five to seven times. The extent of cell lysis was examined using a light microscope and approximately 98 % of amoeboid cells were lysed. The lysates were serially diluted, plated onto Sabouraud agar and incubated at 30 °C for 48 h for colony form unit (CFU) determination. To study the effect of calcium and/or magnesium on the survival of *C. neoformans* during the interaction with amoeba, DPBS was replaced with DPBS containing 0.9 mM calcium and/or 0.5 mM magnesium. To study if the cations affect the adhesion between *C. neoformans* and *A. castellanii*, either or both of *C. neoformans* and *A. castellanii* are pretreated with DPBS containing 0.9 mM calcium and 0.5 mM magnesium for 1 h. Cells were then washed with DPBS. Viability of *A. castellanii* was also determined under the same conditions and time intervals by adding 1:80 dilution of trypan blue stain. The percentage of dead amoeba was determined by counting the number of trypan blue stained cells per total cell number counted. Minimal of 100 cells were counted. Control wells contained *A. castellanii* without *C. neoformans*.

Microscopy and Time-lapse imaging. For measuring the spread area of *A. castellanii* in the presence or absence of calcium and magnesium, *A. castellanii* cells (5 × 105 cells) were allowed to adhere for 2 h on a glass surface of poly-D-lysine coated coverslip bottom MatTek petri dishes with 14 mm microwell (MatTek Corporation, Ashland, MA) in DPBS with or without magnesium and calcium. Images were then taken using a Zeiss Axiovert 200M inverted microscope with 20× phase objective. The area of amoeboid cells (minimum 100 cells) was measured using imageJ software.

For fluorescent imaging, *A. castellanii* cells were washed twice with DPBS and 2 × 10^4^ amoeboid cells were seeded on 8-well chambered coverglass (Nunc, Roskilde, Denmark) in DPBS with or without magnesium and calcium at 25 °C for 2 h. *C. neoformans* were stained with 0.01 % Uvitex 2B (Polysciences, Warminster, PA) for 10 min and wash twice with DPBS. Uvitex 2B-stained *C. neoformans* (2 × 10^4^ cells) were added into *A. castellanii* culture and incubated at 25 °C. After 2 h and 24 h incubation, images were taken using a DAPI filter-equipped Zeiss Axiovert 200M inverted microscope with 20× phase objective. Images were also used for measuring the phagocytosis index which was determined by the number of *C. neoformans* per 100 amoeboid cells.

For time-lapse imaging, *A. castellanii* were seeded (5 × 105 cells) on MatTek petri dishes in DPBS with or without magnesium and calcium. Cells were then incubated at 25 °C for 2 h. Cryptococcal cells (5 × 105 cells/well) were added into amoeba culture. After 15 min incubation to allow fungal cell to settle down, images were taken every 30 s for 24 h using a Zeiss Axiovert 200M inverted microscope with a 10× phase objective in an enclosed chamber under conditions of 25 °C.

The movement of *A. castellanii* were tracked manually using Image J Manual Tracking. Ten amoeboid cells were randomly selected and tracked at the first two hours of incubation with C. neoformans in each tested condition. Total distance, mean velocity and the number of contact between *A. castellanii* and *C. neoformans* were measured. The total distance was defined as the sum of the distance the amoeboid cells travelled from starting point to end point of cell trajectory. The mean velocity was calculated as the mean of all velocity measurements from amoeboid cells moving in each 30 s interval.

### Supernatant toxicity assay

To examine the effect of the secretion of *A. castellanii* on *C. neoformans*, transwell^®^ plates (6-well, polycarbonate membrane, pore size 0.4 µm; Corning) were used to separate *C. neoformans* and *A. castellanii* into two compartments with continuity of saline. *A. castellanii* cells were washed twice and suspended with DPBS containing magnesium and calcium, and 2 × 105 cells were added at the lower compartment. The plates were then incubated at 25 °C for 2 h. *C. neoformans* were washed and suspended with DPBS containing magnesium and calcium, and 2 × 105 cryptococcal cells were added to the inserts. After 0 h and 24 h incubation, *C. neoformans* cells in the inserts were serially diluted, plated onto Sabouraud agar and incubated at 30 °C for 48 h for colony form unit (CFU) determination.

To examine whether the secretion of *A. castellanii* products required direct cell contact between amoeba and fungi, *C. neoformans* were treated with amoeba-conditioned medium collected after incubation of *A. castellanii* and *C. neoformans*. *A. castellanii* cells (1 × 10^4^ cells) in DPBS containing magnesium and calcium were added to 96-well plates and allowed to adhere for 2 h at 25 °C. *C. neoformans* cells were washed twice with DPBS containing magnesium and calcium and 1 × 10^4^ cells were added to wells containing amoeba and the plates were incubated at 25 °C for 24 h. Conditioned media were collected and filtered through 0.22 µm syringe filter. Fresh culture of *C. neoformans* were washed twice with DPBS containing magnesium and calcium, and suspended in the conditioned medium. Fungal cells (1 × 10^4^) were added to wells and incubated at 25 °C. Control wells contain co-incubation of *A. castellanii* and *C. neoformans* as well as *C. neoformans* alone with *A. castellanii* conditioned medium. After 0 h and 24 h incubation, the amoeba were lysed by pulling the culture through a 27-gauge syringe needles five to seven times. The lysates were serially diluted, plated onto Sabouraud agar and incubated at 30 °C for 48 h for colony form unit (CFU) determination.

### Statistical analysis

All comparisons were either analyzed by unpaired two-tailed Student’s t test or two-tailed Fisher’s exact test.

## Acknowledgments

This work was supported in part by 5R01HL059842, 5R01AI033774, 5R37AI033142, and 5R01AI052733.

## Figure legends

**Fig S1.**
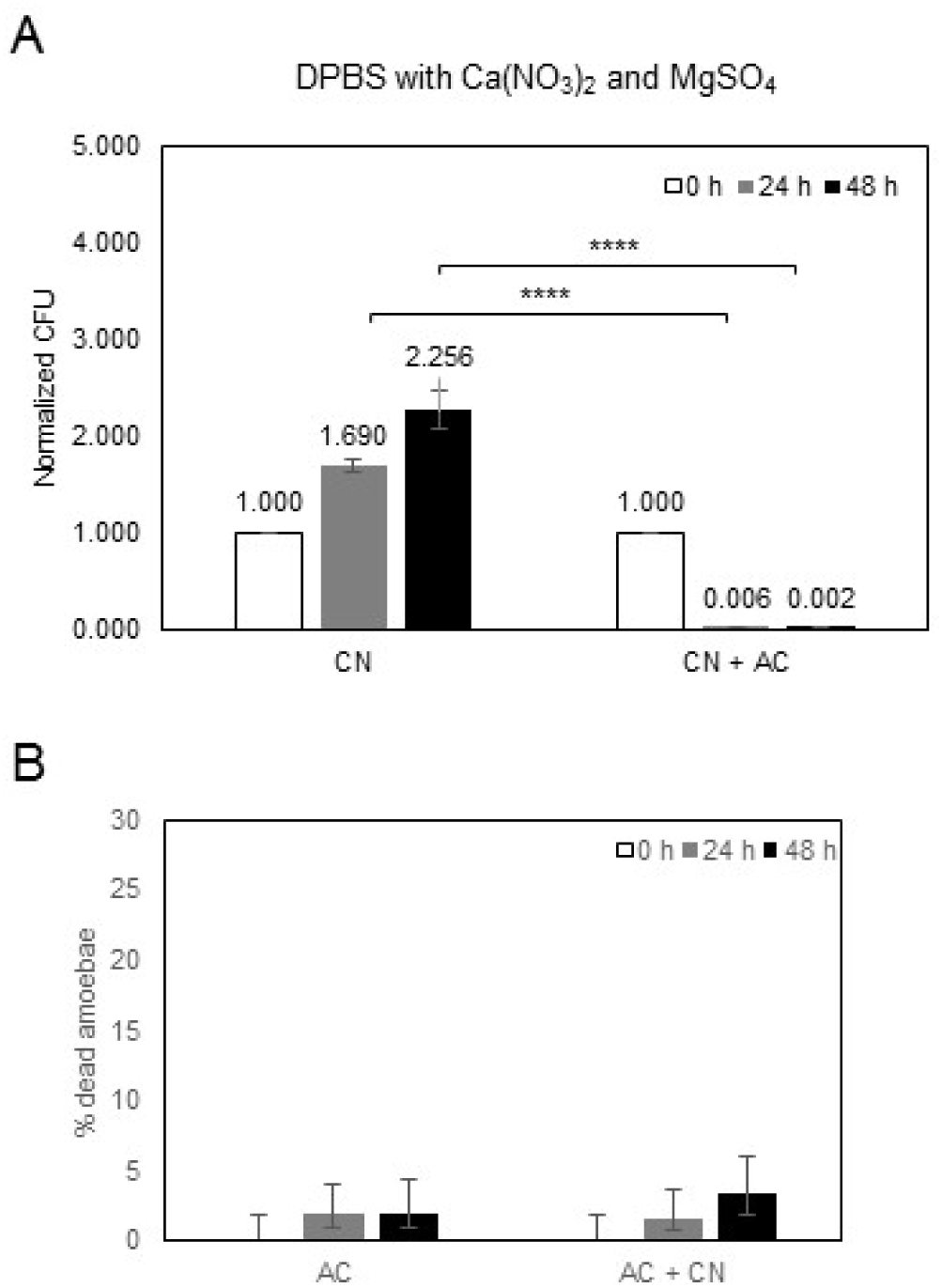
Different forms of magnesium and calcium also affects the outcome of the *A. castellanii*-*C. neoformans* interaction. (A) The assay of *A. castellanii*-*C. neoformans* interaction was performed in DPBS supplemented with magnesium sulfate and calcium nitrate. The survival of *C. neoformans* was determined by CFU after incubation with *A. castellanii* for 0, 24 and 48 h. The CFU counts at 24 h and 48 h were normalized to the initial CFU at time zero. Data represent the mean of three biological samples. Error bars are SD. **** P < 0.0001 by Student’s t test. (B) The viability of *A. castellanii* were determined by trypan blue exclusion assay. The percentage of dead *A. castellanii* is determine by counting the number of blue staining cells per total cell number counted. Three independent biological experiments were performed. Error bars represent 95 % confidence interval of the mean.

**Fig S2.**
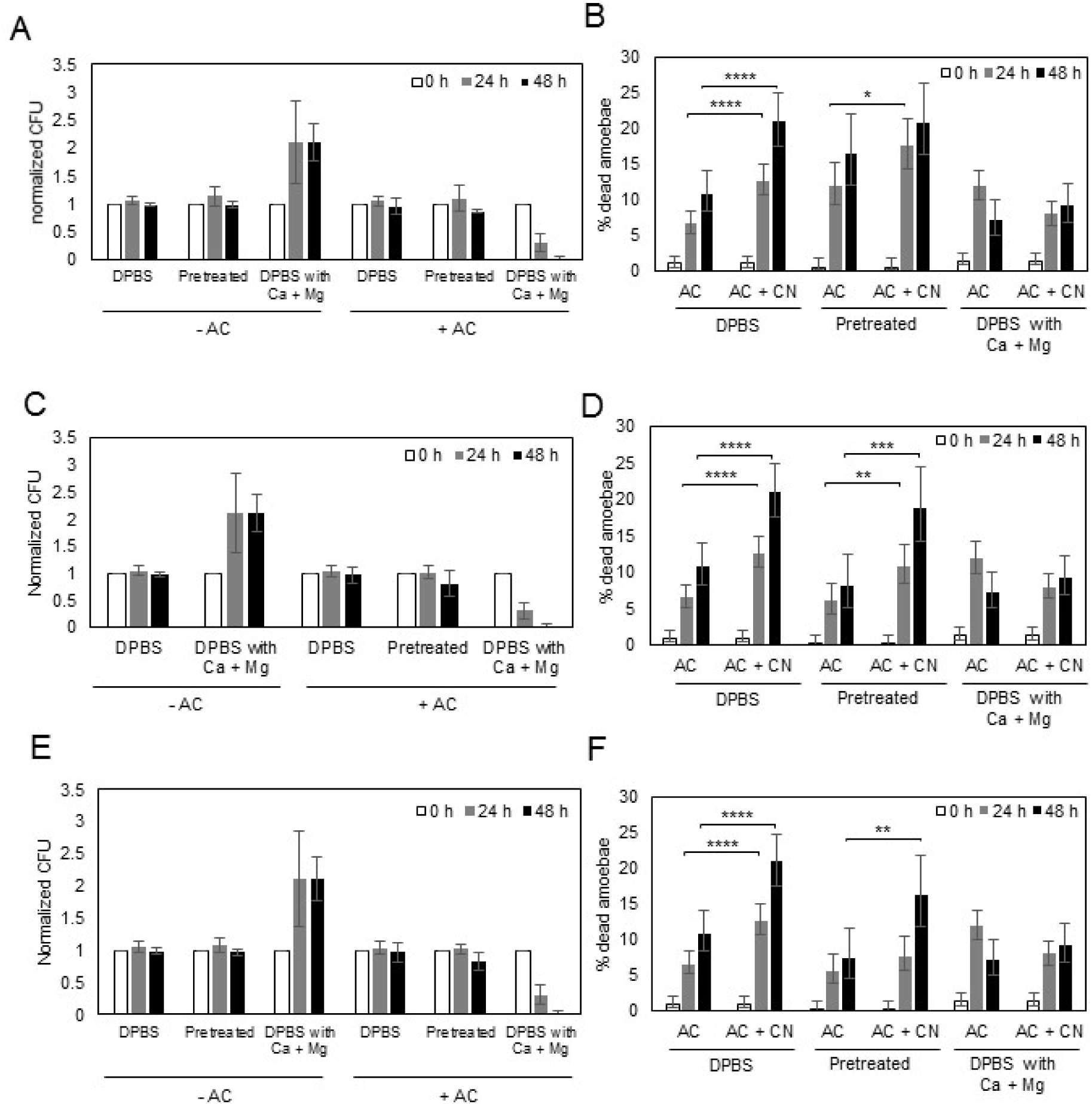
Pretreated *C. neoformans* or/and *A. castellanii* with DPBS containing Ca^2+^ and Mg^2+^ did not affect the outcome of the interaction. The survival of *C. neoformans* and the viability of *A. castellanii* were determined when (A-B) *C. neoformans* was pretreated (C-D), *A. castellanii* was pretreated, and (E-F) both *C. neoformans* and *A. castellanii* were pretreated with DPBS containing Ca^2+^ and Mg^2+^ in prior to the co-incubation of *C. neoformans* and *A. castellanii* in DPBS. Three biological samples were performed in each condition. For the data of the viability of *A. castellanii*, error bars represent 95 % confidence interval of the mean. * P < 0.05 ** P < 0.01 ***P <0.001 **** P < 0.0001 by Fisher’s exact test by Fisher’s exact test. For the data of *C. neoformans* survival, error bars are SD.

Movie S1. Time-lapse microscopy of co-incubation of *C. neoformans* and *A. castellanii* in DPBS with Ca^2+^

